# Decoding Human Emotional Faces in the Dog’s Brain

**DOI:** 10.1101/134080

**Authors:** Raúl Hernández-Pérez, Luis Concha, Laura V. Cuaya

**Author notes:** For correspondence (LVC).

## Abstract

Dogs can interpret emotional human faces (especially the ones expressing happiness), yet the cerebral correlates of this process are unknown. Using functional magnetic resonance imaging (fMRI) we studied eight awake and unrestrained dogs. In Experiment 1 dogs observed happy and neutral human faces, and found increased brain activity when viewing happy human faces in temporal cortex and caudate. In Experiment 2 the dogs were presented with human faces expressing happiness, anger, fear, or sadness. Using the resulting cluster from Experiment 1 we trained a linear support vector machine classifier to discriminate between pairs of emotions and found that it could only discriminate between happiness and the other emotions. Finally, evaluation of the whole-brain fMRI time courses through a similar classifier allowed us to predict the emotion being observed by the dogs. Our results show that human emotions are specifically represented in dogs’ brains, highlighting their importance for inter-species communication.

## Introduction

Emotions provide social cues to interact with the social world. Basic emotions, as expressed through facial expressions (which are universal in humans (***Ekman and Cordaro, 2011***), are automatic physiological responses to certain stimuli, and have a high adaptive value. Moreover, interpretation of emotions in others is fundamental across species for communication between conspecifics (***Anderson and Adolphs, 2014; Leopold and Rhodes, 2010***). Certain species, however, extend their communication necessities beyond their kind. In this sense, dogs are a special species, as they live in a rich social environment that includes humans, making the heterospecific relationship with them a challenge. Their ability to fit their environment is strongly dependent on their capacity to communicate and interpret emotional cues from humans. Dogs seem to be particularly adept at reading human faces, as they are sensitive to human emotional faces, which is evident by their longer visual examination of an emotional face over a neutral face (***Hori et al., 2011***) (but also see (***Racca et al., 2012***)).

A smile is very powerful: it can cheer up our day and even modify our behavior. What is interesting is that we are not the only species capable of recognizing happiness in a human face —dogs can, too. For example, a previous study showed that dogs gazed longer at their owners when they seemed happy (as owners viewed a cheerful movie) than when they seemed sad (***Morisaki et al., 2009***). Another interesting report showed that once dogs learn to discriminate a smiling from a neutral face by viewing photographs of their owners’ faces, they can generalize their learning to other humans of the same gender as their owners (***Nagasawa et al., 2011***). An eye-tracking study (***Somppi et al., 2016***) found that dogs use a conjunction of information from eyes, midface and mouth (i.e., inner face) to process faces in general; however, when looking at a positive human face, dogs spend significantly more time looking at the eyes in comparison to the mouth. Nonetheless, dogs and humans seem to share a similar mechanism to perceive emotional human faces, as they both present a bias to the left hemiface, and they both spend more time looking to the left hemiface of a face with a positive valence (***Racca et al., 2012***).

Moreover, dogs can discriminate other basic emotional expressions in human faces: happiness ((***Albuquerque et al., 2016***), vs. anger (***Müller et al., 2015***), vs. fear (***Merola et al., 2014***), vs. sadness (***Morisaki et al., 2009***), vs. disgust (***Buttelmann and Tomasello, 2013***), and vs. neutral (***Nagasawa et al., 2011; Deputte and Doll, 2011***); anger (***Albuquerque et al., 2016; Müller et al., 2015***), vs. neutral (***Deputte and Doll, 2011***), sadness (vs. happiness (***Morisaki et al., 2009***); fear (vs. happy (***Merola et al., 2014***); and disgust (vs. happiness (***Buttelmann and Tomasello, 2013***). There are no reports about discrimination of surprise. Dogs discriminate emotions deriving information beyond local cues (like the eyes) and can generalize this ability when they are shown novel stimuli (***Müller et al., 2015***). The fact that dogs are sensitive to emotional human faces and capable to discriminate between emotions might explain their remarkable behavioral flexibility in social contexts (***Miklosi et al., 2004***). Beyond dogs’ sensitivity and discrimination of emotional human faces, there is some evidence that dogs can interpret human emotions and modulate their behavior accordingly. For example, dogs choose to first explore a box identified by a human through a facial expression of happiness than a box related to disgust or fear on behalf of the human, suggesting that dogs, to some degree, use human emotional expressions as references (***Merola et al., 2014; Buttelmann and Tomasello, 2013***). Moreover, dogs can correctly identify happiness and anger through only visual or auditory information, showing the cognitive capacity of dogs to extract and integrate emotional information from at least these two sensory modalities (***Albuquerque et al., 2016***).

Behavioral evidence, however, does not provide a clear prediction about the cerebral mechanisms that underlie processing of emotional human faces in dogs. Our goals are to describe the cerebral correlates of the perception of happy human faces in dogs and explore whether a human face expressing an emotion generates a discernible pattern of activity in a dog’s brain using functional magnetic resonance imaging (fMRI). To this end, we used multivariate pattern analysis (MVPA) with a linear support vector machine classifier (LSVM). The main advantage of MVPA is its high sensitivity, and its main assumption is that cognitive states are represented by different patterns of neural activity (***Norman et al., 2006; Kragel and LaBar, 2016***). Our hypotheses are that processing of happy human faces involves the temporal cortex, and that there exist specific neural signatures related to processing these positive visual stimuli that make possible their discrimination to other emotions.

## Results

### Experiment 1

Here, our goal was to describe the cerebral correlates of happy human faces in dogs. We chose happiness because there is more evidence of the perception of this than any other emotion. Eight trained and awake dogs participated. They remained still inside the scanner watching Happy and Neutral human faces in a block design for five runs (the duration of each run was 192.5 s), while fMRI were acquired using blood-oxygen level dependent (BOLD) contrast images (see Materials and Methods).

To identify face-sensitive regions, we analyzed the cerebral activity resulting from viewing both happy and neutral faces in the contrast Faces > Baseline (Figure 1A). The results were overlaid on the Datta atlas (***Datta et al., 2012***). The activity related to viewing faces, regardless of their emotion, includes bilateral occipital cortex and left temporal cortex.

**Figure 1.**
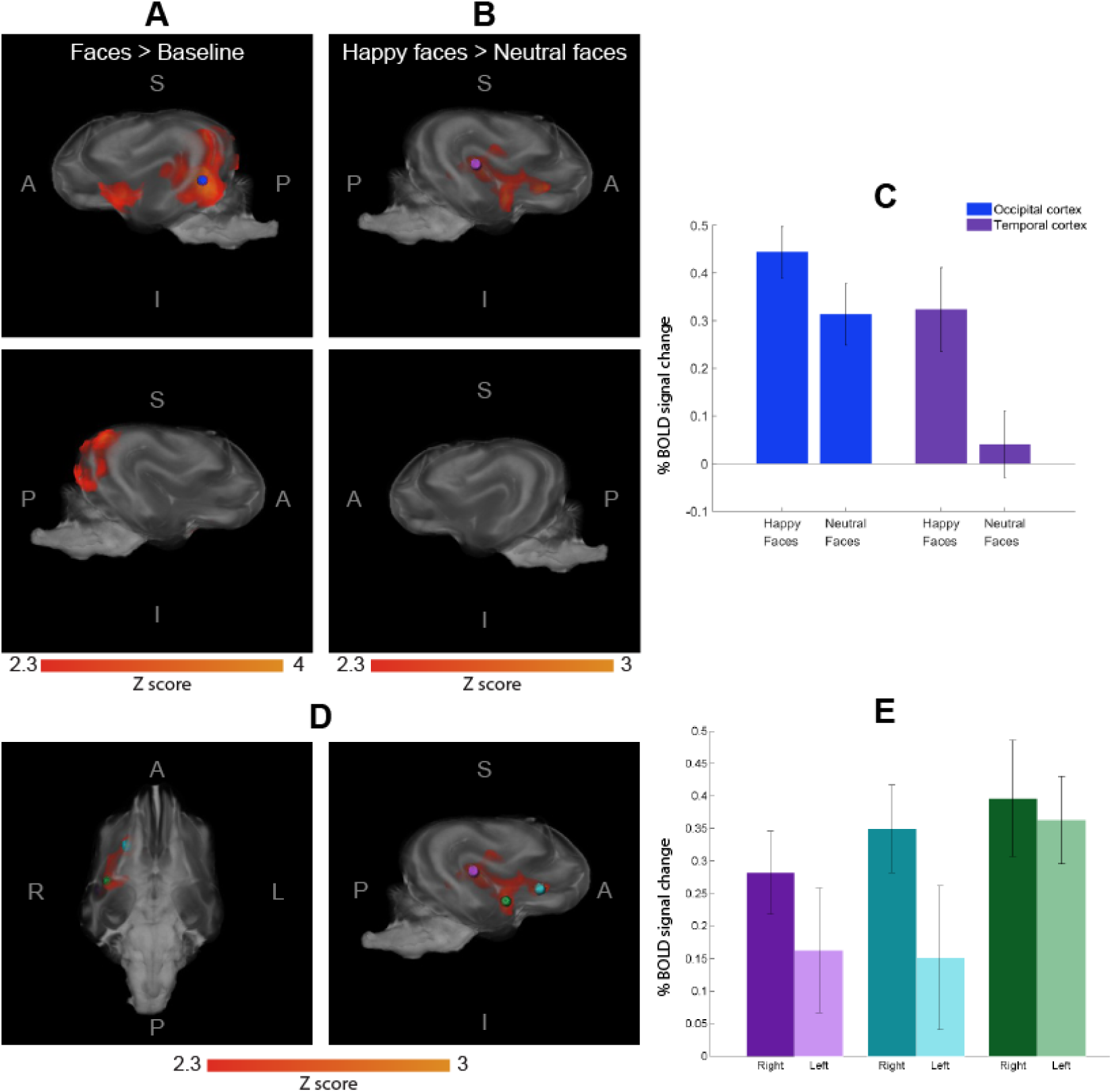
Differences in the cerebral response to happy vs. neutral faces (n = 8). Volume rendering in lateral and inferior views, showing resulting clusters overlaid on the Datta atlas 2012. **A**. Faces > Baseline; the blue sphere indicates the voxel with largest z value located in the occipital cortex. **B**. Happy > Neutral faces; the purple sphere is centered at the voxel with maximum z value, located in the temporal cortex. **C**. BOLD signal change between happy and neutral faces in the spheres of the local maxima in both contrasts. **D**. The three local maxima within the cluster Happy > Neutral faces, all in the right hemisphere. Similar spherical regions were created on the left hemisphere at the same locations. **E**. BOLD signal change between happy and neutral faces derived from these spherical regions (dark and light bars for right and left hemispheres, respectively). Vertical lines represent standard error. S = Superior, I = Inferior, L = Left, R = Right, P = Posterior, and A = Anterior.

To describe the dog’s brain correlates of perception of happy human faces, we contrasted BOLD activity between the two viewing conditions (i.e., Happy human faces > Neutral human faces), which resulted in a large cluster in the right temporal cortex and extend to caudate (25,327 mm^3^, Figure 2B). Figure 2C shows that BOLD signal response towards happy faces and neutral faces is indistinguishable in the occipital cortex, but larger in magnitude in response to viewing happy faces in the temporal cortex. To evaluate inter-hemispheric asymmetries, we extracted the BOLD signal change from spheres of 5 mm radius centered at three local maxima of the resulting right-temporal cluster, located in the Sylvian, Proreus, and Straight Gyri (***Datta et al., 2012***). The difference in BOLD signal change between happy and neutral faces in each sphere is shown to the right in Figure 2E. Contrary to the right-hemispheric BOLD activity, we did not find any significant differences in the activity elicited by happy or neutral faces in the left hemisphere temporal regions (paired t test, p > 0.05). The response in the Straight gyrus is the highest and least lateralized in comparison with the other structures. Coordinates, localization, and z value of local maxima are shown in Table 1. The opposite contrast (Neutral human faces > Happy human faces) showed no significant differences.

**Table 1.**
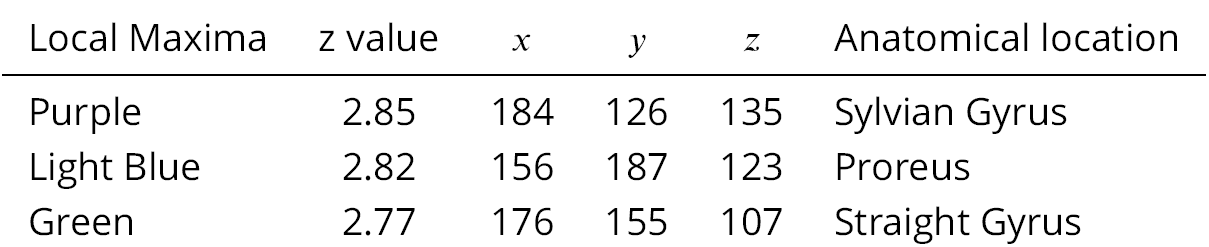
Local Maxima resulting from the contrast Happy Faces *>* Neutral Faces. Coordinates and location are given in mm according to the Datta atlas. Colors are matched with Figure 1.

**Figure 2.**
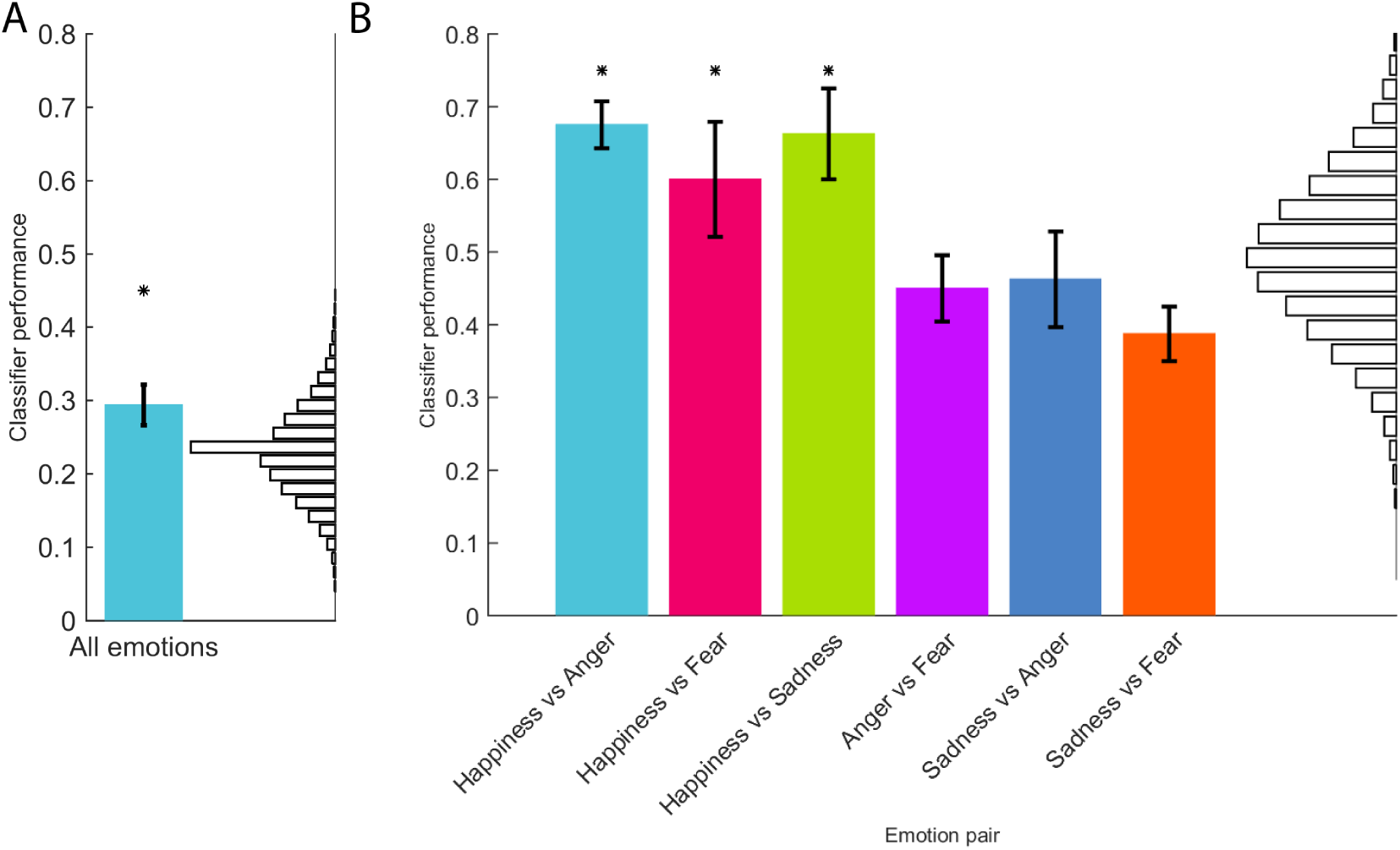
Classifier performance for emotions (n=4). A LSVM classifier was trained using the time series from voxels contained within the cluster identified in Experiment 1 as sensitive to happy human faces. **A**: Comparison of all emotions. **B**: Classifier performance at discriminating between two emotions. Vertical lines denote the standard error. Asterisks represent a significant performance (binomial test p < 0.05). Random classification is shown on right side in black in both panels.

In brief, Experiment 1 provides direct evidence of the cerebral processing of happy human faces in dogs through fMRI. As predicted, we found cerebral activity related to the processing of happy faces in the right temporal cortex.

### Experiment 2

Processing of emotions is not performed in unique cerebral areas, but rather through broad networks in the brain (***Hamann, 2012; Lindquist et al., 2012; Kassam et al., 2013***). It is therefore possible that certain cerebral areas are involved in the processing of multiple emotions, which makes it difficult to study basic emotions with classical univariate approaches. An alternative approach to analyze fMRI data is MVPA, which allows information related to neural representations from distributed patterns in the brain to be decoded. By using MVPA in humans, it has been possible to classify neural signatures associated to specific emotions, and using these patterns it is possible to predict which emotion is being processed in the brain (***Kassam et al., 2013***). In Experiment 2, from the brain activity measured with fMRI in four dogs, we decoded the perception of happiness, sadness, anger and fear using MVPA with a LSVM classifier. We chose to use only these four basic emotions because even for humans it is difficult to discriminate between fear and surprise and between anger and disgust (***Jack et al., 2014***).

#### Experiment 2A

Due to the design of Experiment 1, there is a possibility that the resulting cluster related to processing of happy faces responds to emotional faces in general and not exclusively to happy faces. To address this problem, we further studied four dogs that had participated in Experiment 1 and presented them with new images of emotional human faces while we acquired functional images. Each dog experimented ten runs of stimulation and data acquisition, with each run containing blocks of visual stimulation consisting of faces expressing happiness, sadness, anger or fear; in total each dog experienced 20 blocks of each emotion divided in ten runs.

Using only the time series derived from the cluster related to processing happy faces processing (i.e., result from Experiment 1; Figure 1) we evaluated whether our classifier was able to discern each pair of emotions. From these data, classification performance was above chance (p < 0.05) only when discriminating between happiness and any of the other three emotions (Figure 2).

In summary, independent data was used to demonstrate that the cluster found in Experiment 1 is particularly related to processing of happiness, and not emotions in general. As predicted, viewing happy human faces produces a specific neural signature in the dog’s temporal lobe that makes possible to discriminate it from the activity related to viewing other human emotions.

#### Experiment 2B

With the results from Experiment 2A we could not exclude the possibility that dogs were merely processing emotional valence, as we were only able to discriminate between positive (i.e., happiness) and negative (i.e., anger, sadness and fear) emotions. Experiment 2B was designed to explore the cerebral patterns related to four basic emotions. While the classifier in Experiment 2A was trained with right temporal lobe data only (as this region was identified as being related to processing of happy human faces), this experiment evaluates classifier performance when using activity patterns derived from the entire brain.

We found it was possible to decode the emotion being viewed at the group (Figure 3A) and at the individual level (Figure 3B). We then tested whether it was possible to discriminate between each possible emotion pair, as it was possible that the detection of a single emotion enhanced the predictions. In other words: What if only specific cerebral patterns relative to one or two emotions exist? To test this we performed an analysis only using the cerebral patterns of two emotions each time. The classifier could discriminate above chance level for each emotion pair (Figure 3C) and was able to distinguish even between two negative emotions. Next, we performed a Representation Similarity Analysis (RSA) (***Connolly et al., 2012***), to evaluate the similarity of the different cerebral patterns related to emotions (Figure 3D). This information is represented in a dendrogram (Figure 3E). The most similar emotions were sadness and anger in the first group, happiness in another group and fear in a third group.

**Figure 3.**
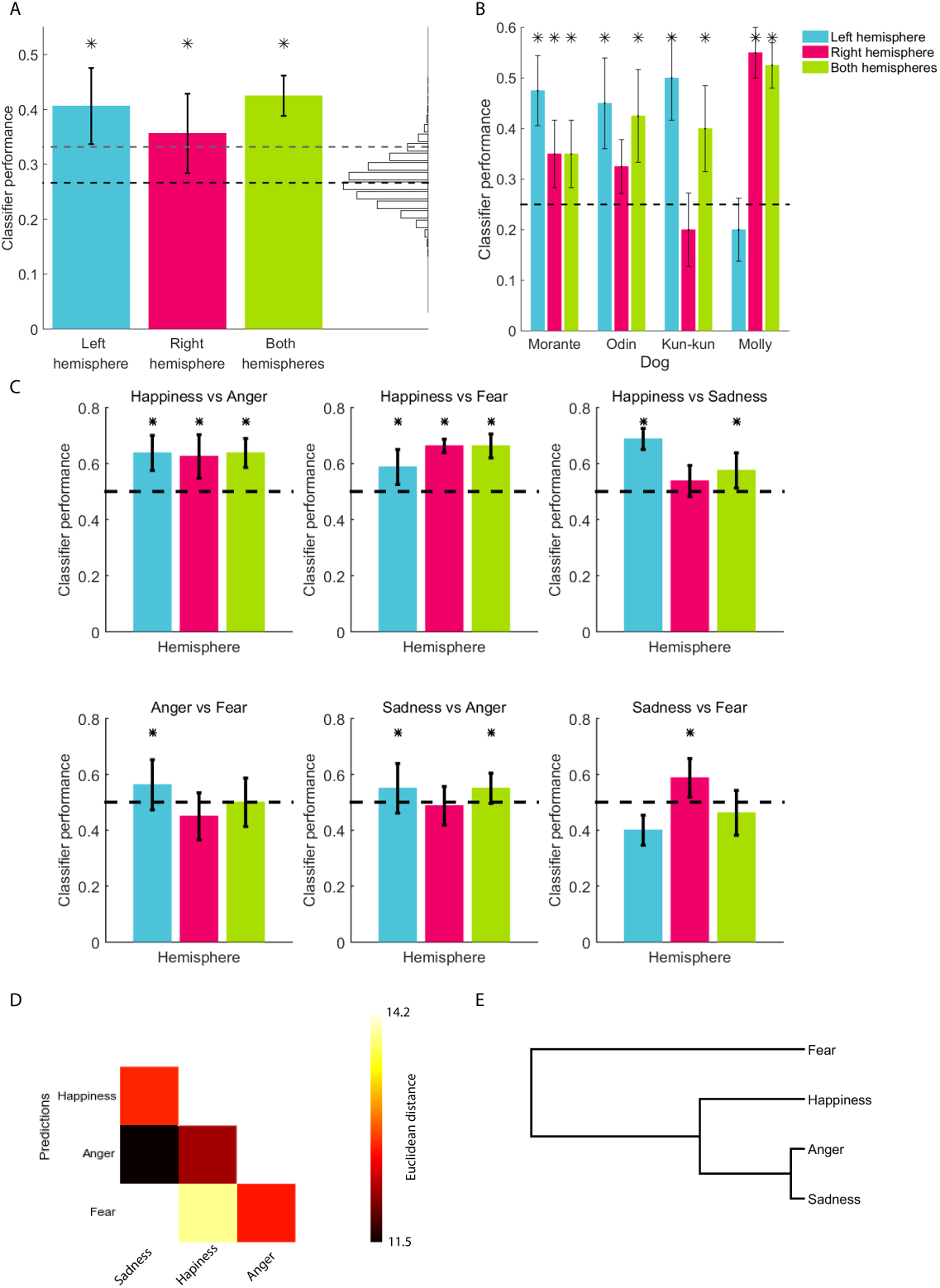
Classification of cerebral patterns in dogs towards emotional faces (n=4). To decode the cerebral patterns we used multivariate pattern analysis. **A-C**: Classifier performance using a leave-one-out cross-validation. Black dashed line indicates chance level; vertical lines denote the standard error; asterisks represent a significant result (binominal test, p < 0.05). A. Group level results. Random classification is shown on right side in black (classifications above the dashed gray line have p < 0.05). **B**: Individual level results. **C**: Predictions by each emotion pair. Representation similarity analysis color-coded in **D** and as a dendrogram in **E**.

The information necessary to decode each emotion is not localized in the temporal cortex. Indeed, in each dog the voxels used to make the prediction were scattered around the brain (Figures 4A and 5). Even taking a different number of voxels the predictions were consistently better when using the information from both hemispheres, and the best predictions for each hemisphere and the full brain plateau at around 50 voxels (Figure 4B). These findings suggest that emotions are represented in a distributed brain network in dogs.

**Figure 4.**
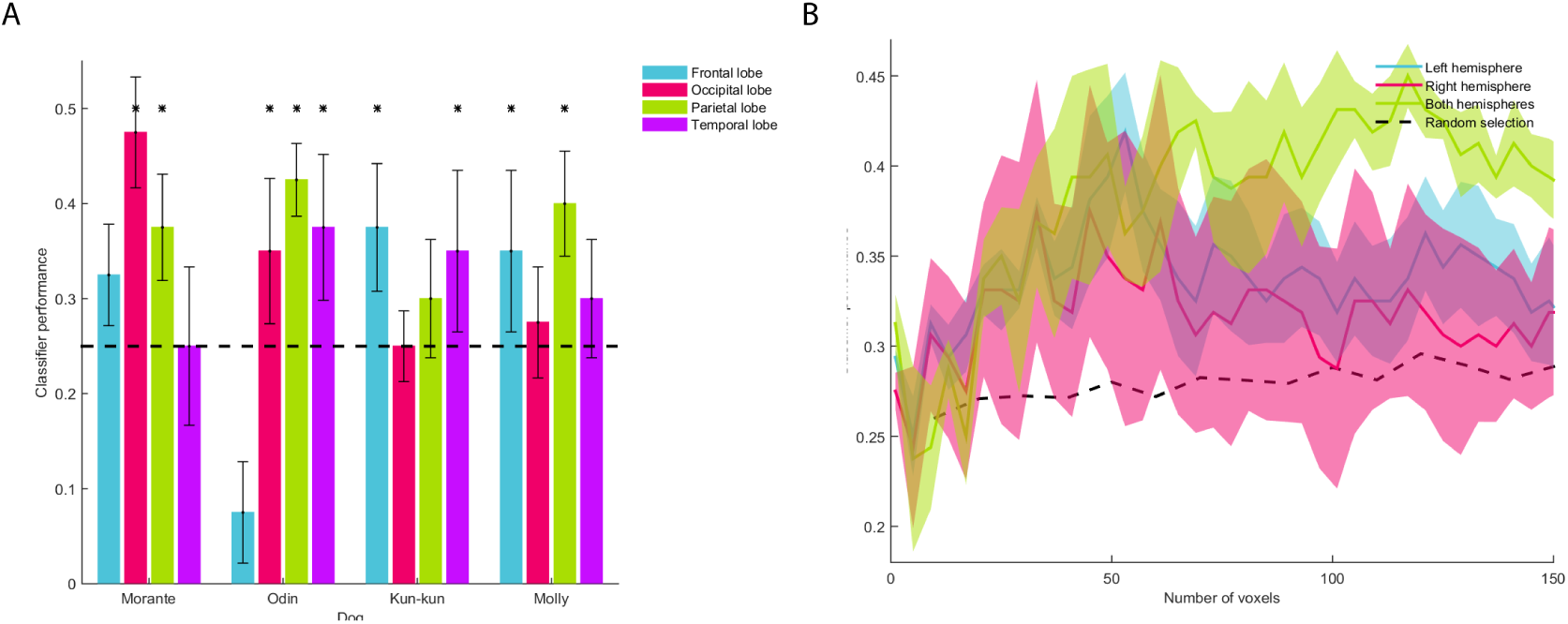
Dogs process the emotions seen in human faces through distributed cerebral networks. **A**. We performed the MVPA in each dog using the information from each brain lobe. The dashed line represents chance level; asterisks represent a significant result (binomial test, p < 0.05). **B**. The classification was performed with an increasing number of voxels. The classifier prediction of randomly selected voxels is represented by the dashed black line. Error bars in A and colored areas in B represent the standard error.

## Discussion

The goals of this study were to describe the cerebral correlates of the perception of happy human faces, and explore whether four basic emotions (happiness, anger, sadness and fear) generate a discriminable pattern of activity in dogs. As predicted, processing of happy human faces involves the temporal cortex, yet other regions (e.g. caudate) are also recruited. Also, we found that happy faces induce a specific signature of cortical activity that makes it possible to discriminate it from other emotions. With our data, it was also possible to discriminate between negative emotions, thus suggesting that dogs not only process emotional valence.

We found cerebral activity related to the processing of happy faces in the right hemisphere, mainly in the temporal cortex (Figure 1). Previous studies found that the perception of human faces (vs. objects) elicits a strong response in the temporal cortex (***Cuaya et al., 2016; Dilks et al., 2015***), and our current study extends said findings, showing that the strongest response to happy faces is located in the superior portion of the temporal lobe. In humans, the superior temporal sulcus (STS) is crucial for social communication, including the dynamic aspects of face perception, such as emotional expressions (***Kanwisher and Yovel, 2006; Haxby et al., 2000; Andrews and Ewbank, 2004; Allison et al., 2000***). We suggest that in dogs the superior temporal cortex could play a similar role in the processing of dynamic aspects present in a happy human face.

The model of the distributed human neural system for face perception (***Haxby et al., 2000***) proposes a hierarchical structure of face perception with a core system and an extended system. The core system includes the occipital and temporal cortex with the STS related to the changeable aspects of faces (like emotional expressions), in contrast to other face-sensitive regions like the occipital cortex, that are not modulated by emotional facial expressions. The processing, or extended, system includes other cortices beyond the occipitotemporal visual extrastriate cortex. Our findings in dogs are analogous to those seen in humans, and compatible with the distributed model for face perception. The occipital lobe was involved in the early stages of face perception, but was not involved in coding of emotions (Figure 1A). Contrastingly, the activity seen in the frontal regions (Proreus) and caudate is evidence of the participation of these structures as part of an extended system for face perception in dogs (Figure 1D). Further supporting this distributed model in dogs the most informative voxels to predict emotions in each dog were broadly spread throughout the brain (Figure 5).

**Figure 5.**
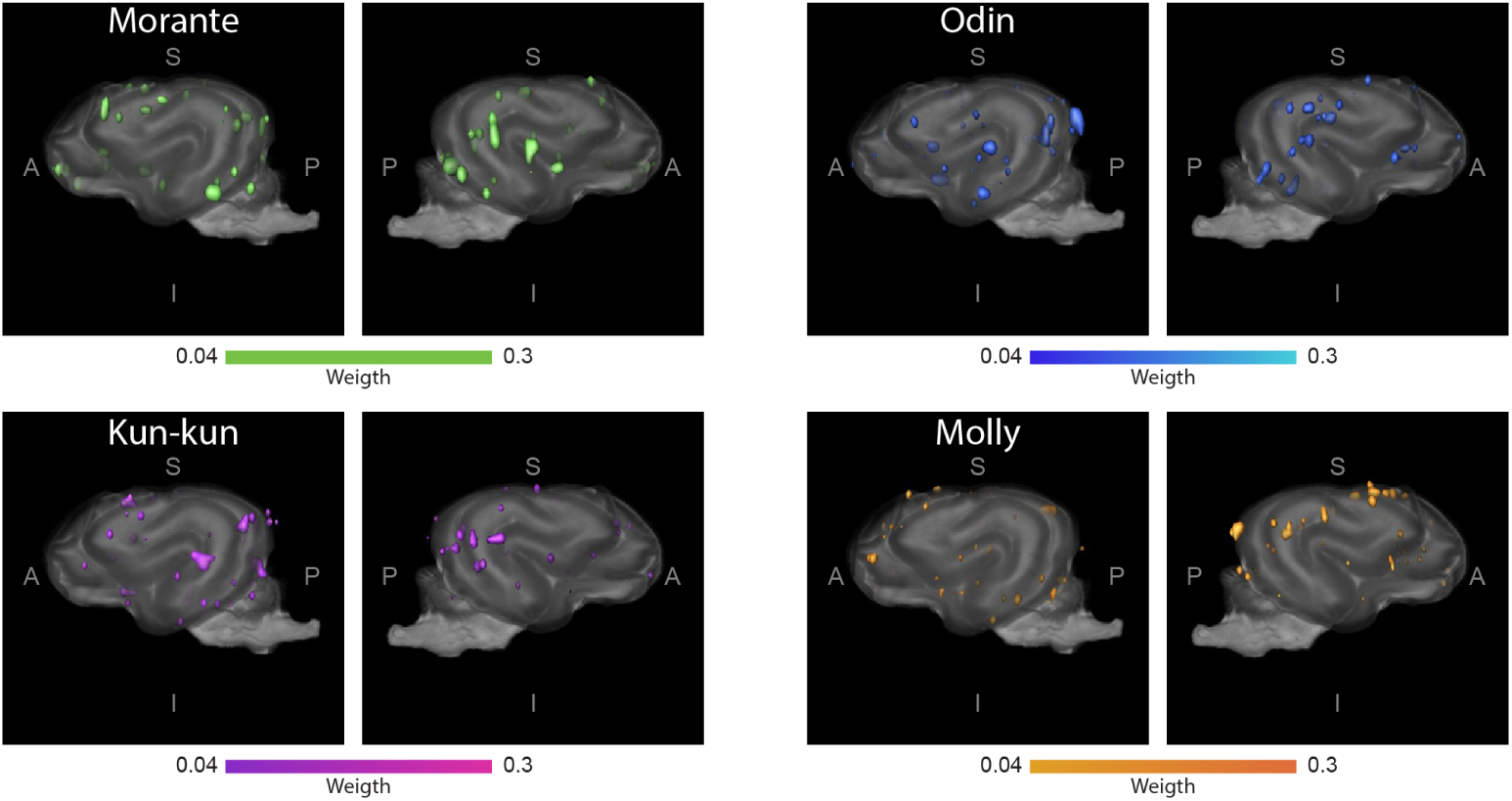
Distributed networks in the dog’s brain used to decode human emotional faces. For each dog, voxels that predict the observed emotion are seen scattered around the brain (overlaid on the Datta atlas 2012). Voxels used for the classification where the top 1% of an ANOVA-based feature selection ranking. The color gradient indicates the weight assigned by the LSVM classifier.

The cluster related to viewing happy human faces included the caudate. Other fMRI studies with dogs have also shown activation of related to reward (***Berns et al., 2013***), familiarity of scents (***Berns et al., 2015***) and neutral human faces (***Cuaya et al., 2016***). In a study in which two groups of dogs were trained to identify either happy or angry faces, both groups of dogs learned to discriminate facial emotions, yet the group trained to identify happy faces learned significantly faster (***Müller et al., 2015***). While the authors of said study suggested that dogs recognized an angry face as an aversive stimulus, the inverse is just as likely, with dogs showing accelerated learning due to the rewarding nature of happy faces. Moreover, a recent neuroimaging study found that the caudate is functionally connected to the auditory regions only when dogs listen to verbal praise with meaning and positive intonation (***Andics et al., 2016***); intonational information alone was not enough to recruit the caudate. Therefore, it is possible that happy human faces not only served as rewarding stimuli for dogs, but also represented meaningful stimuli, consistent with the importance of emotions to dogs, a social species. Besides, happy human faces appear to play a major role in establishing inter-species attachment. In recent years, oxytocin has been highlighted as a key player in this process: dogs spontaneously look longer at their owners when they express positive feelings (***Morisaki et al., 2009***) and this gazing promotes a release of oxytocin in dogs (***Nagasawa et al., 2009, 2015***). This is similar to the enhancing effects of oxytocin on emotion recognition in humans (***Shahrestani et al., 2013; Marsh et al., 2010; Domes et al., 2013***), While we did not directly investigate the role of oxytocin in human face perception, said findings are suggestive of our current results may be associated to this hormone.

It was possible to discriminate between all the pairs of emotions from patterns of brain activity (Figure 3C). However, our classifier showed considerably better performance differentiating happiness from the other three stimuli, as compared to any other pair of emotions. It was possible to correctly identify the emotions being viewed using data from all the brain, or from the right or left hemispheres separately (except vs. sadness in the right hemisphere). We posit that the arousal elicited by fear induces a pattern of brain activity with larger differences than patterns associated to the other emotions, as illustrated in the dendrogram resulting from RSA (Figures 3, D and E). Also, we show that the dog’s brain not only separates the presented emotions by valence, because even negative emotions showed a specific brain patterns that allowed the classifier to discriminate between them. This is a surprising result given that other studies had failed to find behavioral differences between anger vs. sadness and happiness vs. neutral (***Barber et al., 2016***); threatening vs. pleasant and pleasant vs. neutral (***Somppi et al., 2016***); happiness vs. neutral (***Buttelmann and Tomasello, 2013***); fear vs. happiness (with a stranger person) and fear vs. neutral (***Merola et al., 2014***). While behavioral studies suggest that it is necessary to compare opposite emotions to find differences (***Somppi et al., 2016; Müller et al., 2015; Merola et al., 2014; Buttelmann and Tomasello, 2013; Barber et al., 2016; Nagasawa et al., 2011***), we found differences in brain activity related to happy and neutral faces (Figure 1) and it was possible to discriminate all emotion pairs by applying MVPA (Figure 3C). It is intriguing why dogs have specific cerebral patterns towards faces expressing sadness, anger, fear and happiness, yet sometimes they fail to discriminate or use these emotions at the behavioral level. Additional brain processes, environmental cues or other variables may ultimately sway the dog’s behavior.

Our results do not show a clear lateralization. In Experiment 1 all brain activity related to happy faces appeared only in the right hemisphere, yet we believe that this is an effect of the statistical threshold used, because the left hemisphere also shows activity related to happy faces (Figure 1E). In Experiment 2B we didn’t find lateralization either; at the group level both the left and the right hemisphere can discriminate between emotions above chance level (Figure 3A). And at the individual level (Figure 3B) there is not a clear pattern of laterality. The Valence Model (***Honk and Schutter, 2006***) suggests that the negative emotions are processed in the right hemisphere, but this is not supported by our results as the patterns from the right hemisphere can neither discriminate between anger and fear nor between sadness and anger (Figure 3C). Further arguing against the lateralization of the emotion representation, we found that the predictions improve when the classifier takes information from both hemispheres the predictions (Figure 4B).

A strong point in this study is that the faces seen by the dogs are unfamiliar to them and of both genders in Experiment 2. Dogs more readily use emotionally-charged information when expressed by the owner but not by a stranger (***Merola et al., 2014***). Dogs also show difficulty generalizing the discrimination of happiness on faces that are opposite to the gender of their caretaker (***Nagasawa et al., 2011***). In Experiment 2, the classifier accuracy may have been better if we had used facial emotion expressions from their caretakers, but in this exploratory study, we aimed at a more general identification of different emotions. Behavioral studies about emotion perception in dogs have shown differences across breeds (***Mehrkam and Wynne, 2014; Gácsi et al., 2009***) (but also see (***Pongrácz et al., 2005***)) and we are therefore cautious in the extrapolation of our results to other breeds. The majority of our participants were Border Collies, dogs that are better at following social cues than non-working dogs (***Wobber et al., 2009***), spend more time looking at humans and are more willing to interact with them (***Kis et al., 2014***) and show an increased sensitivity to the social effects of oxytocin (***Kovács et al., 2016***). It is possible that Border Collies also have a greater ability to interpret human emotions, and future studies are needed to determine if there are behavioral and cerebral differences in the perception of emotions across breeds. The small sample is other limitation of this study, secondary to the challenges intrinsic to training dogs for the acquisition of functional images. Moreover, imaging sessions had to be kept as short as possible to ensure dogs’ attention and compliance. Also, BOLD signal is inherently noisy in humans but more so in dogs, and it may be crucial to develop coils tailored to the anatomy of dogs (***Huber and Lamm, 2017***). Future studies will be needed to replicate and expand our results.

In conclusion, Experiment 1 provides the first evidence that a happy human face generates a broad activity in the dog’s brain, and implies that dogs can discriminate at the brain level between happy and neutral faces. Along with these results, Experiment 2 showed that dogs have specific patterns of neural activation towards four basic emotion expressions in human faces. The neural signatures for each emotion can be found at the individual level, demonstrating the high sensitivity of dogs’ brains toward human emotions. Our results highlight the importance of human emotions to dogs, since they are a social species adapted to live in an anthropogenic environment.

## Materials and Methods

### Experiment 1

#### Participants

Eight healthy dogs participated in the study. All were medium breeds (six Border Collies, one Labrador, and one Golden Retriever) ranging in age from 18 to 53 months. The sample included four neutered males, one neutered female, and three unneutered females. All dogs were pets, lived with human families, and were well socialized with humans and other dogs. During the study the dogs lived with their human families without changes in their routine aside from the training and imaging protocols described herein. All procedures were performed in compliance with Association for the Study Animal Behavior (ASAB) guidelines, and the Bioethics Committee of the Institute of Neurobiology of the Universidad Nacional Autónoma de México approved the study. The main caretaker of each dog gave informed consent. The dogs were awake and unrestrained during all sessions.

Of the eight dogs included in this study, seven had previously participated in an fMRI experiment (***Cuaya et al., 2016***) and had been trained to remain still inside a scanner in a sphinx position while watching images projected during the acquisition of fMRI. Those dogs did not require additional training. The new participant (Molly) was trained using the same procedure as the other seven dogs (***Cuaya et al., 2016***).

#### Design and Stimuli

We used images of human faces as stimuli and used a block design. There were two types of blocks: faces with neutral expressions and faces with happy expressions. Each block was presented for 7 s with an inter-trial period of 12.25 s used to estimate baseline activity (during which they observed a white screen with a small cross). Each fMRI acquisition (run) had a duration of 192.5 s and included 5 blocks of happy faces and 5 blocks of neutral faces. Blocks were presented in a pseudo randomized order, and the same type of block was never presented two consecutive times. In total, each dog experienced 5 runs (each run had a different order of blocks). Dogs experienced only 1-3 runs with rest periods between them in each imaging session, and different sessions occurred on different days. We decided to use short runs to maximize the attention that dogs pay to the stimuli and minimize the possibility of a dog falling asleep. Thus, fMRI acquisition was completed throughout 2-3 non-consecutive days for each dog. Visual stimulus presentation was controlled by PsychoPy2 (***Peirce, 2007***). The lights in the MRI suite were turned off during paradigm presentation, in all sessions an experimenter was present in MRI suite to ensure the dog’s compliance during acquisitions, and the stimuli were projected onto a screen in front of the dogs at a distance of 1.5 m.

Four different photographs of faces composed each block. All images were extracted from the AR Face Database (***Martinez, 1998***), so all faces were unfamiliar to the dogs. We presented the same persons with neutral and happy expressions. Each image was presented only once during each run to avoid habituation to the images. In order to have stimuli as natural as possible, all images were in color, presented in a frontal view, and the size of the projected images was similar to that of a real face (15 cm x 20 cm), as this size has been used in other studies (***Cuaya et al., 2016; Huber et al., 2013; Nagasawa et al., 2011***). The gender of the faces presented to each participant was the same as the main caretaker: four dogs (one male dog and three female dogs) saw female faces and the other four participants saw male faces. We decided to show a specific gender to each dog, as dogs show lower accuracy in a behavioral task when probed with an unfamiliar face of a person of the opposite gender of their main caretaker (***Nagasawa et al., 2011***).

#### Data acquisition

All images were acquired at the National Laboratory for Magnetic Resonance Imaging in the Institute of Neurobiology of the Universidad Nacional Autónoma de México. We used a 3 T Philips Achieva TX scanner and a two-channel SENSE Flex Small coil, which was attached to the dogs with Velcro. To help the dogs maintain the sphinx position, we used a chin rest to support their heads. Dogs were fitted with ear muffs to provide noise protection during scanning. During the acquisition one experimenter remained inside the scanner room (out of the dog’s sight) and visually monitored the dogs to make sure they were awake and attentive. The dogs could leave the session at any time.

For anatomical reference we also acquired a T1-weighted structural image with a turbo spin echo sequence with 1 × 1 × 1 mm^3^ resolution covering the whole brain with 75 slices. Blood-oxygen-level dependent (BOLD) images covered the whole brain and were acquired with a gradient-echo echo-planar imaging (EPI) sequence (28 coronal slices, 3 mm thickness, no gap; TR = 1.75 s; TE = 30 ms; 2ip angle = 90°; FOV = 224 × 240 mm^2^; acquisition matrix 112 × 120; spatial resolution 2 × 2 × 3 mm^3^; 110 volumes and 5 dummy scans).

#### Image Analysis

The brain was extracted from each run using manual segmentation. All functional analyses were done with FSL (***Jenkinson et al., 2012***) version 4.19. Images were preprocessed for temporal correction, and spatial smoothing was performed using a Gaussian kernel with FWHM = 5 mm. Motion correction was performed using MCFLIRT, and runs showing motion greater to head rotation than 1° or translation greater than 3 mm were discarded (less than 10% of runs; if a run was discarded, a new acquisition was obtained until each dog had five good-quality runs). As maintenance of the sphinx position requires muscle tone, loss of which would result in major head movement, the criteria used for motion correction of imaging data further guaranteed that the dogs remained awake. Functional images were spatially normalized to an anatomical image of each dog and to a digital atlas of the dog’s brain (***Datta et al., 2012***).

We used the General Linear Model for the statistical analyses, including the stimuli vectors of happy and neutral faces as regressors. Each run was analyzed individually in a first level. Regressors were convolved with the canonical hemodynamic response function modeled as a Gamma function. The five runs of each participant were first analyzed using a fixed-effects analysis. A random-effects analysis was used to study common activations within the eight participants. To describe the cerebral regions involved in the perception of happy faces, we analyzed the contrast happy faces vs. neutral faces in the entire brain. The resulting statistical parametric maps were corrected for multiple comparisons using random field theory (***Worsley, 2001***) (cluster-forming threshold z > 2.3 and p_cluster_ < 0.05). To localize and label the cerebral structures and report the coordinates, we used the digital atlas of Datta et al. 2012. We created a sphere of 5 mm radius around the voxel with the local maxima that resulted from the contrast happy faces > neutral faces to extract the BOLD signal change.

### Experiment 2

#### Participants

We studied four healthy and neutered (ages 26 to 48 months, 3 males) Border Collies, all whom had participated in the Experiment 1. We followed the same guidelines of Experiment 1.

#### Design and Stimuli

The stimuli were human faces that expressed happiness, sadness, fear and anger. All stimuli were extracted from the Karolinska Directed Emotional Faces database (***Lundqvist et al., 1998***). We used a block design (each block was presented for 7 s and composed of 4 different images of the same emotion, followed by a baseline of 12.25 s) in each run. An emotion block was presented twice in a pseudo-random order and all the faces belonged to a single gender, which was switched between scans. The total duration per run was 166.25 s. Each dog experienced ten runs in total and a maximum of two runs per day. The projection and dogs’ position were the same as in Experiment 1.

#### Data acquisition

We used the same procedure that in Experiment 1, the only difference being that each run included 95 volumes.

#### Image Analysis

For MVPA, preprocessing was performed using FSL (***Jenkinson et al., 2012***). Images were motion-corrected to the first volume of the first run of each participant. Images were brain-extracted and temporally realigned. The images were neither smoothed nor filtered. We used PyMVPA software package (***Hanke et al., 2009***) and the LibSVM’s implementation of the linear support vector machine (LSVM). Each acquisition was linearly detrended, Z-scored and time shifted 6 s to account for the peak of the hemodynamic response; volumes corresponding to the baseline period were removed. Volumes corresponding to each of the block types were averaged. We followed a leave-one-out cross-validation scheme in which the runs were split into two sets: a training set composed of 9 runs and a test set composed of the remaining run. A LSVM classifier was trained using the training set and tested using the test set; this procedure was repeated using each run as a training set and the others as a training set. For a given training and testing fold, all categories were used, and the classifier performed a 4-way classification in which only a classification that matched the emotion was considered as correct and any other as incorrect. To obtain the distribution of our data under a random classification condition (Figures 2 and 3A, right side), we performed a permutation test in which we randomly swapped the block labels, trained, tested the classifier and repeated the procedure 10,000 times for each mask and each dog. The resulting distributions were compared using a Kolmogorov-Smirnoff test, which showed no significant difference between the distribution (p > 0.05). The three distributions are collapsed in Figure 3. The RSA (***Connolly et al., 2012***) represents each emotion as a vector in a high-dimensional space in which each voxel response represents a dimension. A Euclidean distance is calculated from the resulting vectors. These indicate how similar two emotions are in their cerebral representation.

The general linear model was used as implemented in FSL in a similar way as described in Experiment 1. Each emotion (i.e., happiness, anger, fear, sadness) was used as a regressor and convolved with the canonical hemodynamic response function modeled as a Gamma function. Each run was analyzed individually in a first level; posteriorly, the ten runs for each participant were used for single subject fixed-effect analyses. Finally a random-effect analysis was used for group-level inferences. Statistical parametric maps were thresholded using random field theory (cluster corrected z > 2.3 and p < 0.05).

## Acknowledgments

We are deeply grateful to our participants and their caregivers for their cooperation, patience and friendship: Andrea Dávila and Lourdes Guajardo (Zilla), Ariadna Ríos and Ariel Mendoza (Hera), Daniel Ramírez (Kora), Lenin Ochoa and Jessica Moreno (Morris), Luis Nájera and Liza Guerrero (Molly and Morante), and of course to Odín and Kun-kun. We thank The Dog Project Family for providing their anatomical masks used in Experiment 2b. We also thank Erick Pasaye and Juan Ortiz, and the staff at the National Laboratory for magnetic resonance imaging, for their support during image acquisition. We are grateful to Jessica Gonzalez-Norris for proofreading and editing and to Azalea Reyes and Fernando Barrios for her helpful comments. Raúl Hernández and Laura Cuaya are doctoral students from Programa de Doctorado en Ciencias Biomédicas, Universidad Nacional Autónoma de México (UNAM) and received fellowships 260381 and 260395, respectively, from CONACYT. This study was partially funded by Conacyt (IE252-120295 181508) and UNAM-DGAPA (I1202811 and IN212811).

